# Uric acid accumulation in cockroach wings

**DOI:** 10.1101/2025.11.03.686272

**Authors:** Tomohito Noda, Minoru Moriyama, Toshiyuki Harumoto, Tatsuya Katsuno, Takema Fukatsu

## Abstract

Cockroaches, embracing some 4,500 species in the world, are found in a variety of environments including human settlements. The ecological success of cockroaches is attributable to the ability to endure long periods of starvation and to thrive on nutritionally poor diets, which is underpinned by the capability of storing their own nitrogenous waste products as uric acid in the fat body and also by the endosymbiosis with bacteriocyte-dwelling bacteria *Blattabacterium* that recycle the stored uric acid for synthesis of amino acids. Previous histological studies on cockroaches described that uric acid is accumulated within the fat body, especially in urocytes. Here we report that cockroaches also accumulate uric acid in the wings. In our observation of the German cockroach *Blattella germanica* under different nutritional conditions, we found that well-fed cockroaches exhibited whitish aggregates in the wing veins, which is ascribed to accumulation of uric acid granules. The uric acid granules in the wings disappeared by starvation and were restored by feeding in *B. germanica*. The uric acid granules in the wings were found not only in *B. germanica* but also in other diverse cockroach species.

## INTRODUCTION

Most insects excrete nitrogenous wastes mainly in the form of uric acid via Malpighian tubules (Wigglesworth 1972). In the terrestrial lifestyle of small-size arthropods, production and excretion of uric acid are beneficial for avoiding toxic effects of ammonia by conversion into poorly water-soluble uric acid and thereby for saving water consumption for excretion (Weihrauch and O’Donnell 2021). In addition to excretion, uric acid is involved in various aspects of insect physiology such as nitrogen storage and recycling (Sabree et al. 2009; Patiño-Navarrete et al. 2014), pigmentation and UV protection (Tamura and Akai 1999; Fujii and Banno 2019), antioxidant defense (Matsuo and Ishikawa 1999; Hu et al. 2013; Tasaki et al. 2017), and others. Considering that synthesis of uric acid is energetically costly, adoption of uric acid as the main nitrogenous waste product must reflect its importance in insect physiology and ecology.

Previous studies reported that, in some insects, uric acid is synthesized and stored within fat body cells as solid deposits or in solution, as a means of “nitrogenous waste storage”, when immediate excretion is impossible or unnecessary (Wigglesworth 1972). In cockroaches, mosquitoes, lacewings and others, large quantities of uric acid accumulate in the fat bodies, sometimes forming crystalline deposits (Srivastava and Gupta 1961; Spiegler 1962; Wigglesworth 1987a; Wigglesworth 1987b). In lepidopteran larvae like silkworms, large amounts of uric acid are synthesized by epidermal cells and excreted into the integument, which forms a whitish body color and confers protection against UV damage (Tamura and Akai 1999; Fujii and Banno 2019).

Cockroaches (Insecta: Blattodea) are found commonly in human settlements across the globe and are notorious as hygienic and nuisance pests. Of approximately 4,500 cockroach species, less than 30 species are regarded as pests, and the vast majority have little or no relevance to human activities (Schal and Hamilton 1990; Bell et al. 2007). Instead, cockroaches have adapted to and conquered a variety of environments and ecological niches, such as forests, meadows, deserts, and of course, most recently, anthropic environments (Grimaldi and Engel 2005; Bell et al. 2007). The ecological success of cockroaches may be attributable to their ability to endure long periods of starvation (Abbott 1926; Willis and Lewis 1957; Cornwell 1968) and also their ability to utilize a variety of and often nutritionally poor food sources such as dead wood, detritus and soil (Roth 1980; Mullins and Cochran 1987). A possible contributing factor to their adaptability and resilience is their ability to store and recycle their own nitrogenous waste products, by which they can survive on foods of high carbon-nitrogen ratios. The nitrogen recycling ability of cockroaches is underpinned not only by their capability of storing nitrogen in the form of uric acid in the fat bodies but also by their symbiotic relationship with a bacterial endosymbiont called *Blattabacterium* (Sabree et al. 2009; Lopez-Sanchez et al. 2009). The fat bodies of cockroaches are constituted by three types of cells, adipocytes mainly storing lipids and carbohydrates, urocytes specialized for accumulating uric acid, and bacteriocytes for harboring *Blattabacterium*.

Cockroaches are capable of converting nitrogenous waste products into uric acid, which are mainly stored in urocytes (Mullins and Cochran 1972, 1974, 1976; Cochran et al. 1979; Cochran 1985; Park et al. 2013). *Blattabacterium* is thought to play a role in the catabolism of uric acid and the utilization of the uric acid-derived nitrogenous products for synthesis and provisioning of essential amino acids to the cockroach host (Sabree et al. 2009; Lopez-Sanchez et al. 2009; Patiño-Navarrete et al. 2014).

Thus far, a number of histological studies on the bacteriocytes, the urocytes and the endosymbiotic bacteria *Blattabacterium* have been conducted, which ranged from classic hand-drawn sketches to modern light and electron microscopic observations (Blochmann 1887; Wheeler 1889; Gier 1936; Koch 1949; Bush and Chapman 1961; Buchner 1965; Walker 1965; Milburn 1966; Cochran et al. 1979; Sacchi et al. 1993, 1996, 1998, 2000; Laudani et al. 1995; Lambiase et al. 1997; Park et al. 2013; Noda et al. 2020, 2024). In these previous studies, researchers described uric acid accumulation within fat body cells, especially in urocytes. However, during our research on the cockroach-*Blattabacterium* endosymbiotic relationship under different nutritional conditions, we found that well-fed cockroaches (i.e. under nutritionally rich conditions) exhibit whitish wing veins with some aggregates therein, and demonstrated that it is ascribed to accumulation of uric acid granules. In this study, we report (i) distribution patterns of uric acid granules in the wings of the German cockroach *Blattella germanica*, (ii) histological observations and comparisons of uric acid granules in wings and fat bodies of *B. germanica*, (iii) effects of starvation on uric acid granules in wings of *B. germanica*, (iv) confirmatory identification of uric acid granules of *B. germanica* by liquid chromatography and tandem mass spectrometry (LC-MS/MS), and (v) survey of uric acid granules in the wings of diverse cockroach species.

## MATERIALS AND METHODS

### Insect materials

A stock population of *B. germanica*, derived from around 100 individuals purchased from Sumika Technoservice Corporation, Takarazuka, Japan, was maintained in our laboratory at the National Institute of Advanced Industrial Science and Technology (AIST), Tsukuba, Japan. All insects were reared in plastic containers at 27°C under a 12 h light and 12 h dark regime in a climate chamber (MLR-352H-PJ, Panasonic) with insect feed (Insect Diet I, Oriental Yeast Co., Ltd.) and water. The forest cockroach *Blattella nipponica* and the smokybrown cockroach *Fortiblatta fuliginosa* (also known under the synonym *Periplaneta fuliginosa*) were collected by setting pitfall traps in the AIST campus, Tsukuba, Japan, using dog food as bait and Vaseline to prevent escape. The Turkestan cockroach *Periplaneta lateralis* (also known under the synonyms *Shelfordella lateralis* and *Blatta lateralis*) and the Dubia roach *Blaptica dubia* were purchased at pet stores and were reared similarly to *B. germanica*.

### Dissection and observation of wings and fat bodies

Forewings and hindwings of live insects were cut using dissection scissors, and briefly washed in phosphate buffered saline (PBS: 0.8% NaCl, 0.02% KCl, 0.115% Na_2_HPO_4_, 0.02% KH_2_PO_4_, pH 7.4) and then in 70% ethanol. Cockroaches have two types of fat bodies: the visceral fat bodies that contain adipocytes, urocytes and bacteriocytes, and the peripheral fat bodies that contain no bacteriocytes (Noda et al. 2020). Both the visceral and peripheral fat bodies were dissected in PBS. These samples were mounted in 90% glycerol and observed using either a stereo microscope (M165 FC, Leica) or an upright microscope (DM6 B, Leica).

### Electron microscopy

For backscattered electron scanning electron microscopy (BSE-SEM), visceral and peripheral fat bodies were dissected from adult insects of *B. germanica* in cold PBS. The samples were fixed in 2% glutaraldehyde (Sigma-Aldrich, G5882) in PBS at 4°C overnight, incubated with 2% osmium tetroxide in distilled water at 4°C for 2 h, dehydrated in a graded ethanol series (50%, 60%, 70%, 80%, 90%, 95%, 99%, and 100%), and embedded in epoxy resin (Luveak-812, Nacalai Tesque). The hardened Epon blocks were sectioned using an ultramicrotome (ARTOS 3D, Leica) equipped with a diamond knife (SYM jumbo, 45 degrees, SYNTEK) to obtain 240 nm serial sections. The serial sections were collected on cleaned silicon wafer strips held by a micromanipulator (MN-153, NARISHIGE). The sections were stained at room temperature with 2% (w/v) aqueous uranyl acetate for 20 min and Reynolds’ lead citrate for 3 min. Images were obtained by a scanning electron microscope (JSM-7900F, JEOL).

### Starvation experiment

From the stock population of *B. germanica*, 20 adult insects were randomly selected and subjected to a nitrogen-deficient condition for two weeks, during which the insects were not given any feed but only 10% (w/v) glucose solution. After the starvation treatment, 10 randomly chosen insects were subjected to wing dissection and microscopic inspection. The remaining 10 insects were then returned to their normal feed for a week, after which their wings were dissected for observation. All the other rearing conditions were the same as the stock population.

### Solubilization and LC-MS/MS analysis of uric acid granules

Dissected fat bodies and wings were treated in 1 ml of acetone-hexane (1:1) on a shaker for 30 min to remove lipids and carotenoids, and then incubated in 1 ml of 28% ammonia water on a shaker overnight to dissolve uric acid granules. After a 100-fold dilution with water, the extracts were analyzed using a liquid chromatography-mass spectrometry (LC-MS) system consisting of an ultrahigh-performance liquid chromatograph (Acquity UPLC H-Class, Waters) coupled to a quadrupole-time-of-flight mass spectrometer with an electrospray ionization source (Xevo-G2XS qTOF, Waters). A 1 µL of diluted sample was injected and separated on an ACQUITY UPLC BEH C18 column (2 mm i.d. x 100 mm length, Waters) under a water-acetonitrile gradient at a flow rate of 0.25 mL/min. Positive ions were monitored in the MSe acquisition mode, and fragment spectra were reconstructed from the time frames corresponding to the precursor peak of uric acid at m/z=169.

## RESULTS

### Appearance of whitish aggregates in the wings of *B. germanica*

When we observed well-fed adult males and females of *B. germanica*, whitish aggregates were found in the veins of both forewings and hindwings. In the forewings, the aggregates were mainly found around the basal costal-subcostal region, particularly along the longitudinal and cross veins. In the hindwings, the aggregates were also distributed in the basal costal-subcostal region. There were no notable differences in the distribution patterns of the aggregates between males and females (Fig. 1A-J).

**Fig. 1.**
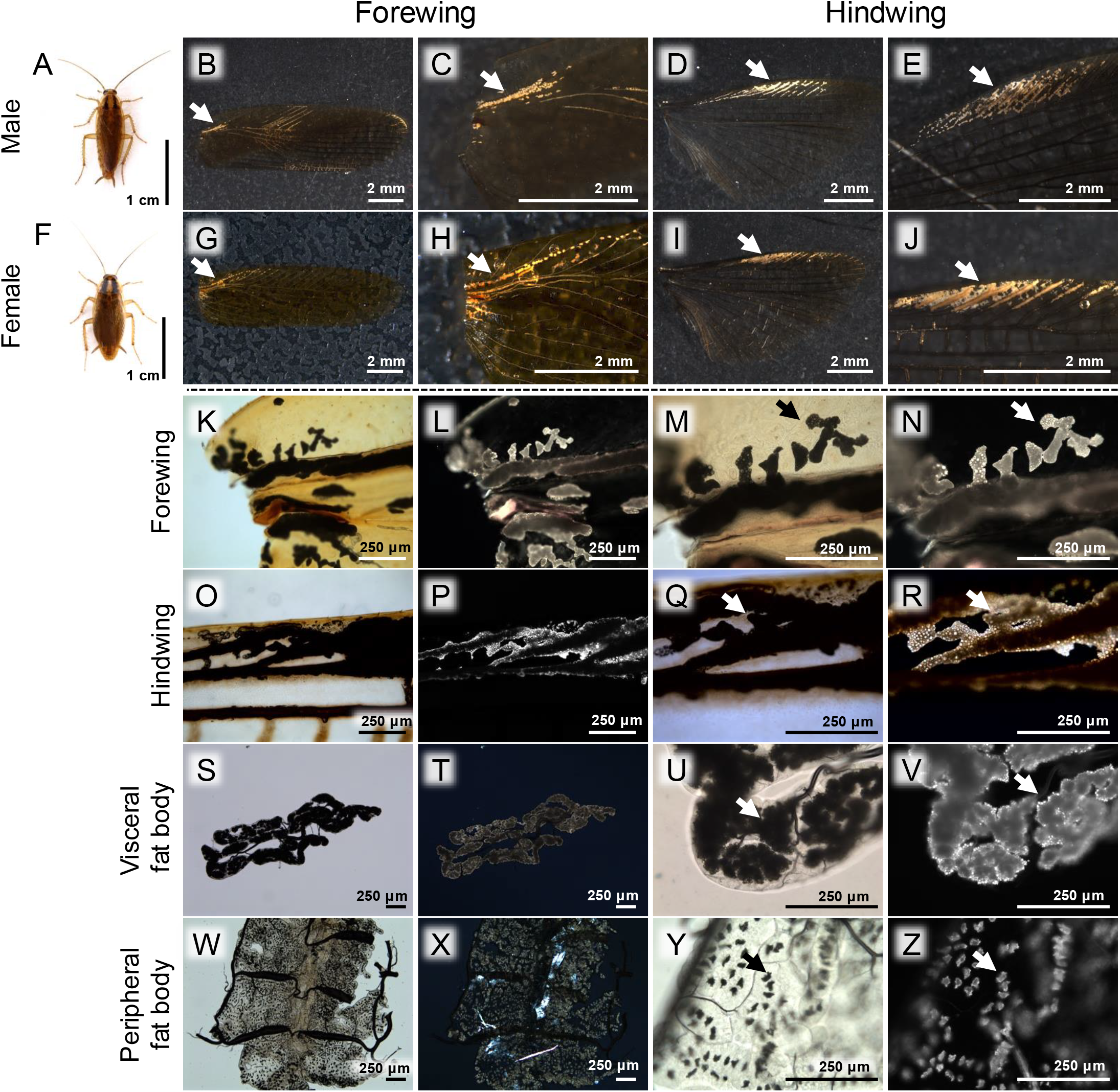
**A–J** Gross appearance of whitish granules accumulated in the wings of the German cockroach *B. germanica*. **A–E** Data obtained from well-fed adult males. **F–J** Data obtained from well-fed adult females. **A, F** Appearance of an adult insect. **B, G** Image of the whole forewing. **C, H** Enlarged image of the granules in the forewing. **D, I** Image of the whole hindwing. **E, J** Enlarged image of the granules in the hindwing. **K–Z** Polarizing microscopic observation of the granules in wings and fat bodies of *B. germanica*. **K–N**, Forewing. **O-R** Hindwing. **S-V**, Visceral fat body. **W-Z**, Peripheral fat body. **K, O, S, W** Transmitted light microscopic image, low magnification. **L, P, T, X** Polarizing microscopic image, low magnification. **M, Q, U, Y** Transmitted light microscopic image, high magnification. **N, R, V, Z** Polarizing microscopic image, high magnification. Arrows point to the granules.

### Polarizing microscopic observation of glittery granules located in wing veins and fat bodies of *B. germanica*

Polarizing microscopic observations of the forewings, the hindwings, the visceral fat bodies and the peripheral fat bodies of *B. germanica* revealed the presence of glittery granules. In the wings, the granules were observed within amorphous scattered cell masses located in wing veins (Fig. 1K-R). In the fat bodies, the granules exhibited characteristic localizations that presumably reflected the locations of the urocytes (Fig. 1S-Z). The accumulation of the granules was more prominent in the visceral fat bodies than in the peripheral fat bodies (Fig. 1U, V, Y, Z; Fig. S1).

### Identification of the granules as uric acid

Uric acid is known for being poorly soluble in water, but it dissolves readily in basic solutions such as ammonia water. When the cockroach tissues were treated with 28% ammonia water, the granules disappeared in all the samples, which became opaque in appearance (Fig. 2A-H). The ammonia water extracts were then subjected to LC-MS/MS analysis, which consistently detected a major peak corresponding to uric acid (Fig. 2I-M, Fig. S2), confirming that the granules are uric acid deposits.

**Fig. 2.**
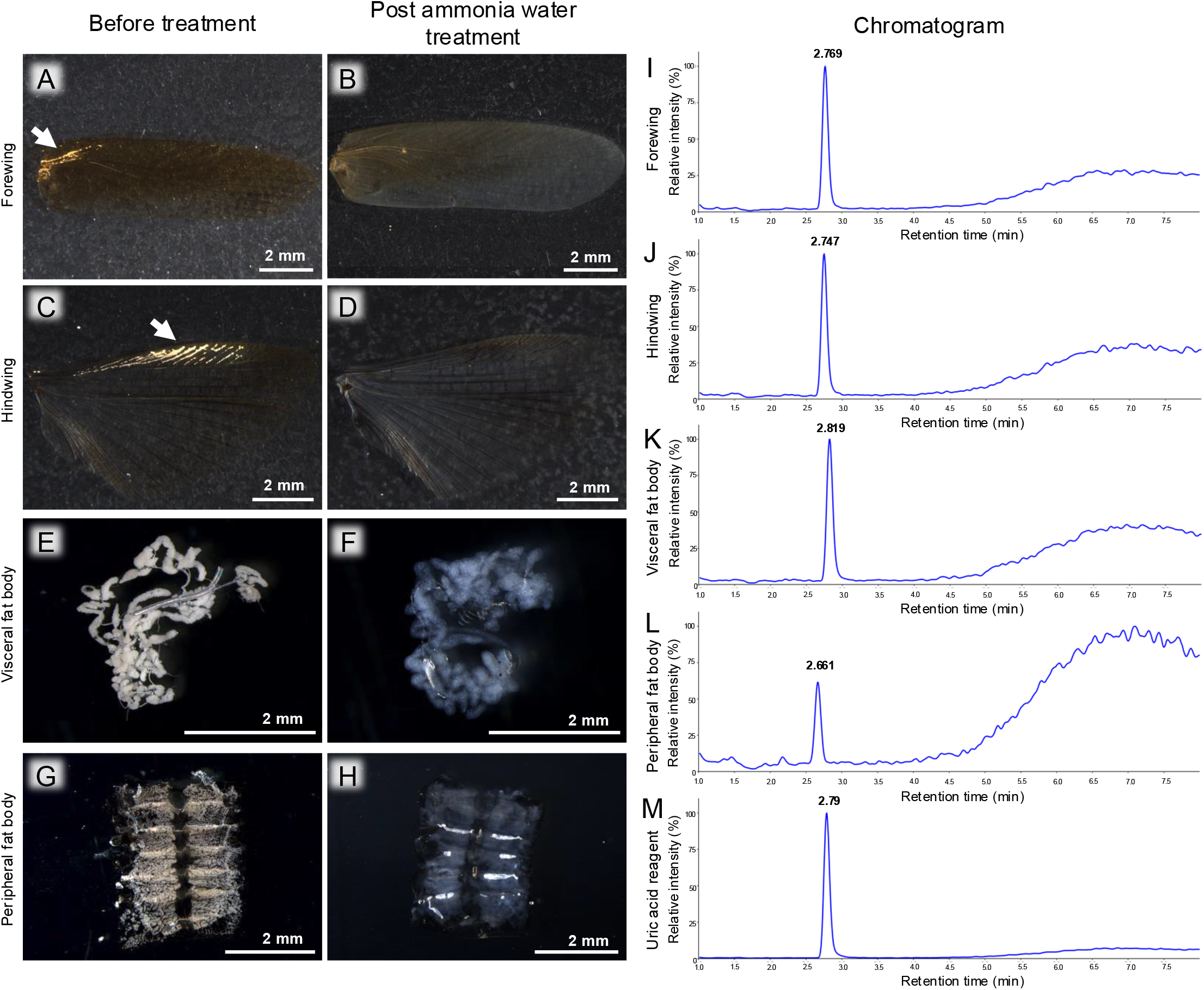
**A–H** Solubilization of the granules from wings and fat bodies of *B. germanica* by ammonia water. **A, B** Forewing. **C, D** Hindwing. **E, F** Visceral fat body. **G, H** Peripheral fat body. **A, C, E, G** Before treatment. **B, D, F, H** After ammonia water treatment. Arrows indicate accumulated granules in wings. **I– M** LC-MC detection of uric acid (m/z=169) extracted from wings and fat bodies of *B. germanica* by ammonia water. **I** Forewing. **J** Hindwing. **K** Visceral fat body. **L** Peripheral fat body. **M** Uric acid reagent (control).

### Effects of starvation on the uric acid granules in wings

Does the nutritional condition of the insect affect the uric acid granules in the wings? To test this, we subjected the insects to nitrogen-deficient conditions by providing only sugar water for two weeks. This treatment resulted in disappearance of the uric acid granules in both the forewings and hindwings of all 10 insects inspected (Fig. 3A-D). Once the insects were returned to regular feed, which is rich in nitrogen, for a week, the uric acid granules recovered in both the forewings and the hindwings in all the insects inspected (Fig. 3E-H).

**Fig. 3.**
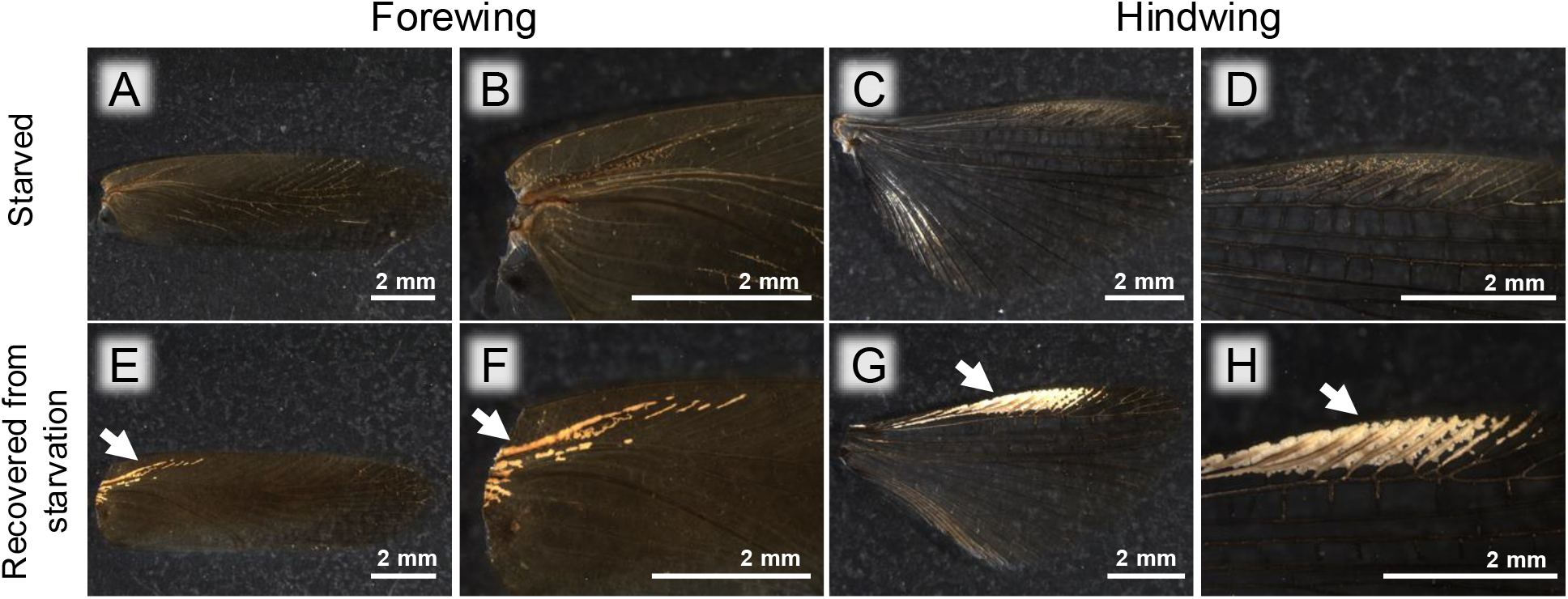
Appearance of the wings of *B. germanica* after starvation and recovery. **A–D** Wings of starved insects fed with only sugar water for two weeks. **E–H** Wings of recovered insects fed with regular feed for a week after the starvation regime. **A, E** Image of the whole forewing. **B, F** Enlarged image of the forewing. **C, G** Image of the whole hindwing. **D, H** Enlarged image of the hindwing. Arrows point to the accumulated uric acid granules.

### Presence of uric acid granules in the wings of diverse cockroach species

Finally, we observed the wings of four other cockroach species, *Blattella nipponica* (Ectobiidae), *Fortiblatta fuliginosa* (Blattidae), *Periplaneta lateralis* (Blattidae) and *Blaptica dubia* (Blaberidae). In all four species, we observed similar granules in both the forewings and hindwings (Fig. 4). The localization of these granules was consistent with uric acid granules observed in *B. germanica*.

**Fig. 4.**
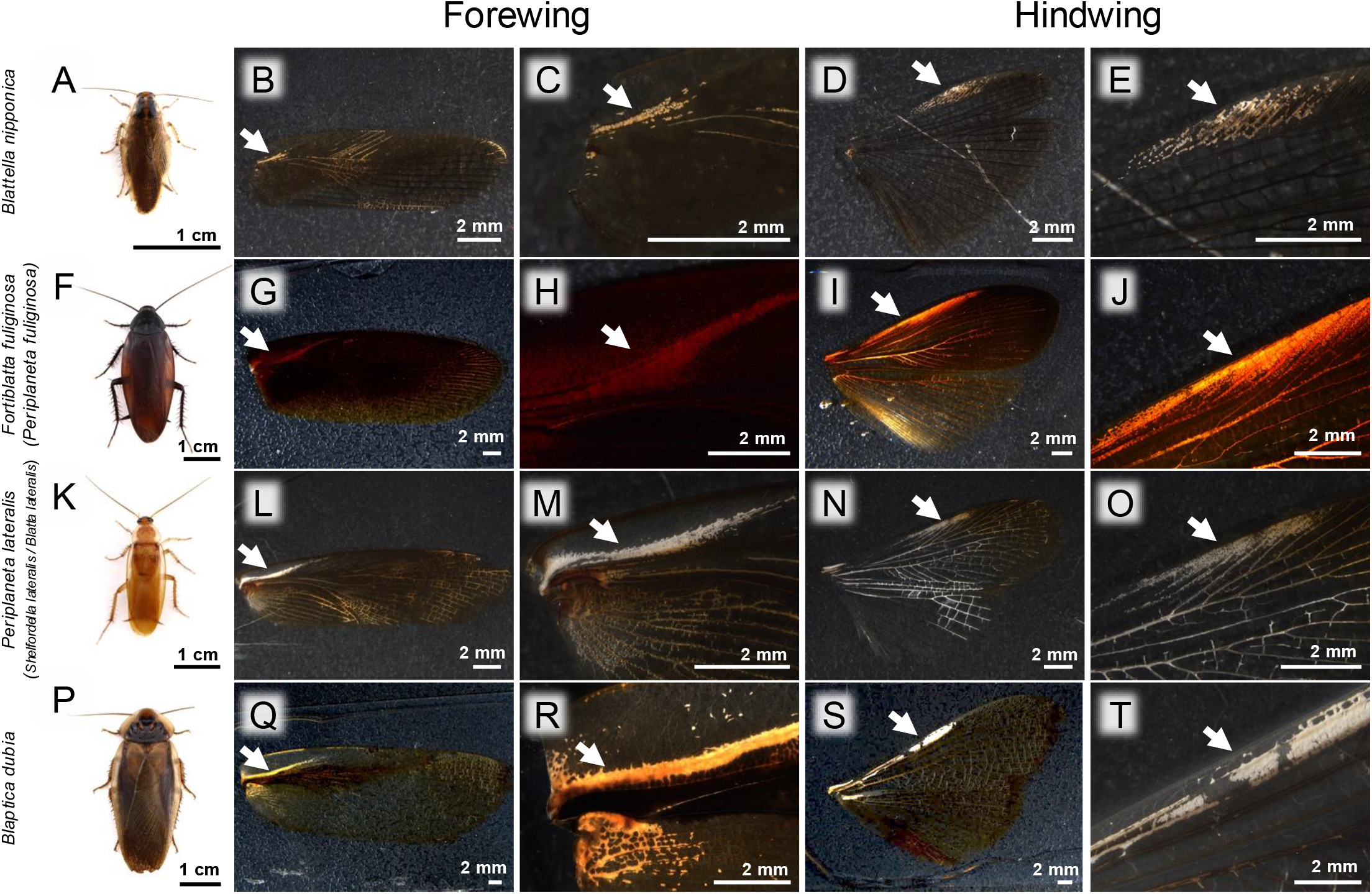
Accumulation of uric acid granules in the wings of diverse cockroach species. **A–E** *Blattella nipponica*. **F–J** *Fortiblatta fuliginosa*. **K–O** *Periplaneta lateralis*. **P–T** *Blaptica dubia*. **A, F, K, P** Appearance of an adult insect. **B, G, L, Q** Image of the whole forewing. **C, H, M, R** Enlarged image of the forewing. **D, I, N, S** Image of the whole hindwing. **E, J, O, T** Enlarged image of the hindwing. Arrows indicate the regions where uric acid granules accumulate.

## DISCUSSION

In previous studies, accumulation and storage of uric acid in fat bodies, particularly within urocytes, have been described (Cochran et al. 1979; Cochran 1985; Park et al. 2013), which were also confirmed by our own histological observations of *B. germanica*. In this study, we found that, in well-fed cockroaches, uric acid accumulates not only within fat bodies in the body cavity but also within other types of cells in the wing veins. The nature of the cells that contain uric acid granules in the wing veins is currently unknown. Possible candidates may be a special type of fat body cells, oenocytes, hemocytes, etc. Future studies using molecular markers will be needed to address this issue.

In some lepidopterans like the silkworm, synthesis and accumulation of uric acid by epithelial cells have been reported, which form its body color and confer defense against UV damage (Tamura and Akai 1999; Fujii and Banno 2019). However, given that the uric acid granules in cockroach wings disappear in response to nitrogen-deficient diets, it seems unlikely to contribute to body coloration or UV protection, but should instead be considered in the context of “nitrogenous waste storage”.

Accumulation of uric acid granules in the wing veins of cockroaches seems counter-intuitive, as it may interfere with the flight function of wings. The biological roles of the uric acid granules in the wings remain elusive. We suppose that the presence of uric acid granules in the wings may entail little biological significance, possibly being a by-product of uric acid accumulation under nutritionally favorable conditions. Meanwhile, the low flight ability of cockroaches and the fact that their wings are neither structurally nor functionally streamlined for flight may allow the existence of substantial hemocoel in the wing veins, which may have enabled the diverse cockroaches to accumulate uric acid granules even in the wings.

## ACKNOWLEDGMENTS

We thank Haruyasu Kohda and Keiko Okamoto-Furuta for their support in electron microscopic analysis, and Genta Okude for his advice on how to remove potential contaminants from the wings. This study was supported by the Japan Science and Technology Agency (JST) ERATO Grant to T.F., M.M., and T.H. (JPMJER1902), the Japan Society for the Promotion of Science (JSPS) Grant to T.H. (JP24H02294 and JP24K08935), and the Hakubi Project of Kyoto University to T.H. The Japan Society for the Promotion of Science (JSPS) Research Fellowships for Young Scientists supported T.N. (JP22KJ1191 and JP21J20814).

## COMPETING INTERESTS

The authors have no competing interests to declare.

## AUTHOR CONTRIBUTIONS

TN conceptualized and designed the study. TN preformed most of the experiments and sample preparation for LC-MS/MS and electron microscopy. MM conducted LC-MS/MS analysis. TH and TK conducted electron microscopy. TN and TF wrote the paper.

## FIGURE LEGENDS

**Fig. S1.**
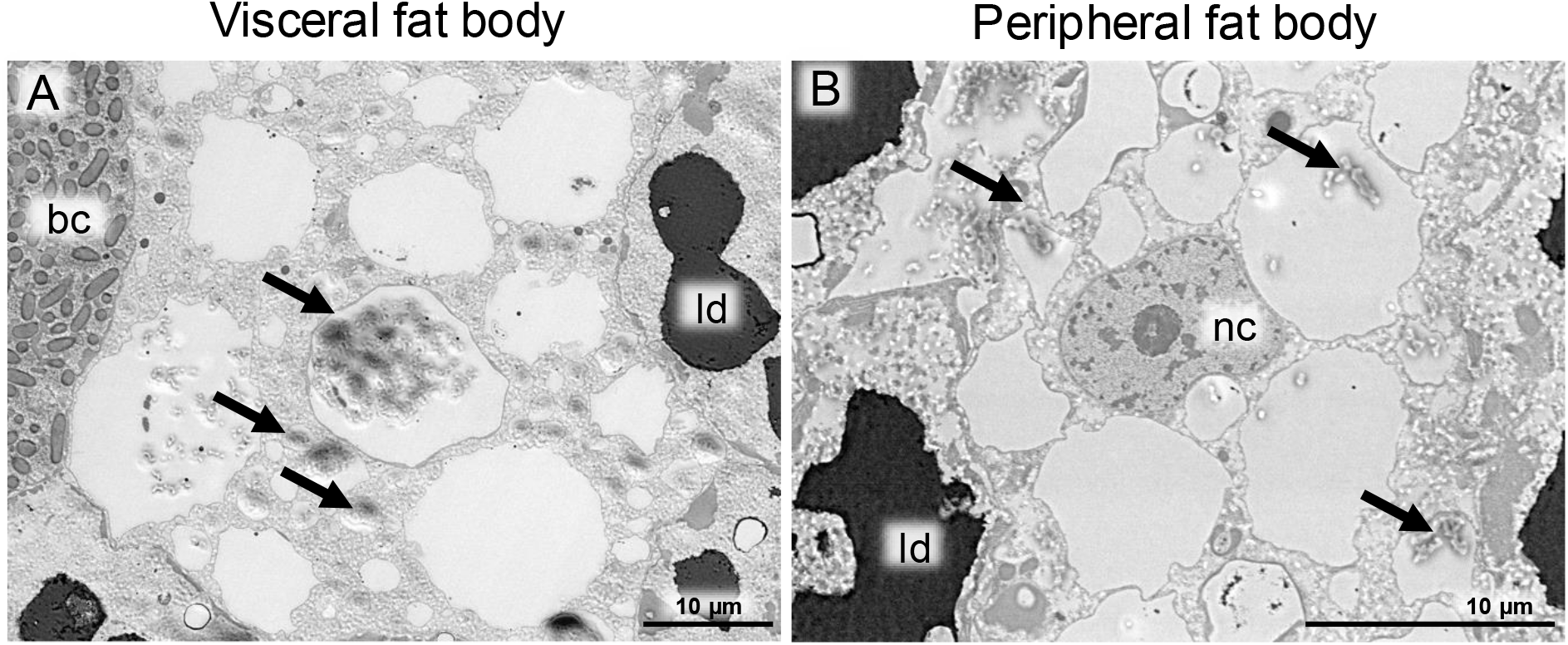
BSE-SEM images of urocytes in fat bodies of *B. germanica*. **A** Visceral fat body. **B** Peripheral fat body. Arrows indicate remaining uric acid deposits. Abbreviations: bc, bacteriocyte; ld, lipid droplet; nc, nucleus.

**Fig. S2.**
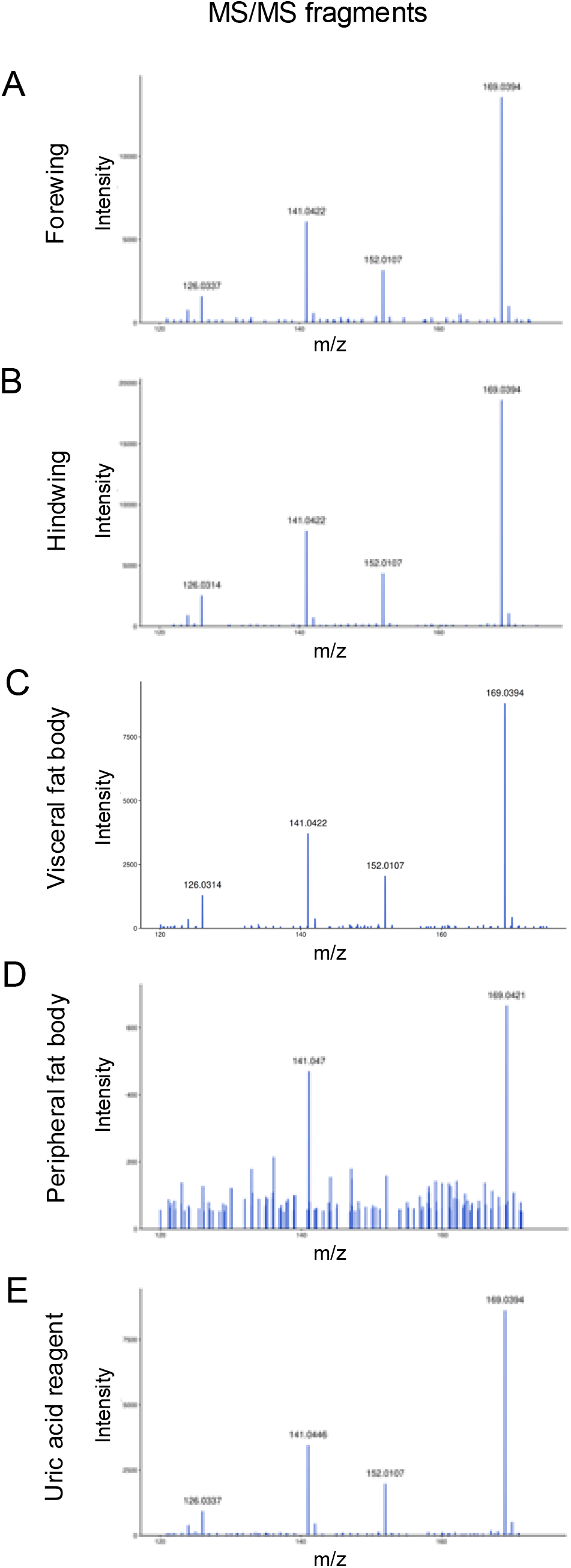
LC-MS/MS identification of uric acid extracted from dissected wings and fat bodies of *B. germanica*. **A** Forewing. **B** Hindwing. **C** Visceral fat body. **D** Peripheral fat body. **E** Uric acid reagent (control).

## Notes

**Statements and Declarations** There are no conflicts of interest.

### Competing Interest Statement

The authors have declared no competing interest.

